# Ectosome effect on endothelial monolayers in hyperglycemic and normoglycemic conditions

**DOI:** 10.1101/2021.11.29.470234

**Authors:** Anna Drożdż, Tomasz Kołodziej, Sonia Wróbel, Krzysztof Misztal, Marta Targosz-Korecka, Marek Drab, Robert Jach, Małgorzata Przybyło, Zenon Rajfur, Ewa Ł Stępień

**Affiliations:** Department of Medical Physics, Marian Smoluchowski Institute of Physics, Faculty of Physics, Astronomy and Applied Computer Science, Jagiellonian University in Kraków, 11 Łojasiewicza St., Kraków, Poland; Department of Molecular and Interfacial Biophysics, Marian Smoluchowski Institute of Physics, Faculty of Physics, Astronomy and Applied Computer Science, Jagiellonian University in Kraków, 11 Łojasiewicza St., Kraków, Poland; Division of Computational Mathematics, Faculty of Mathematics and Computer Science, Jagiellonian University in Kraków, 6 Łojasiewicza St., Kraków, Poland; Department of Nanostructures and Nanotechnology, Marian Smoluchowski Institute of Physics, Faculty of Physics, Astronomy and Applied Computer Science, Jagiellonian University in Kraków, 11 Łojasiewicza St., Kraków, Poland; USI, Unit of Nano-Structural Bio-Interactions, Hirszfeld Institute of Immunology and Experimental Therapy, Polish Academy of Sciences, 12 Weigla St., 53-114 Wroclaw, Poland; Department of Gynecological Endocrinology, Faculty of Medicine, Jagiellonian University Medical College in Kraków, ul. M. Kopernika 23, Kraków, Poland; Department of Glycoconjugate Biochemistry, Institute of Zoology and Biomedical Research, Jagiellonian University in Kraków, 9 Gronostajowa St., Kraków, Poland

**Keywords:** cellular motion, diabetes, endothelial dysfunction, extracellular vesicles, hyperglycemia, wound healing

## Abstract

Extracellular vesicles, namely those larger ones - Ectosomes (Ect), are thought to be important cell-to-cell communication medium. Ect are considered as a potential therapeutic for type-1 and type-2 diabetes mellitus. Ect can be internalized by endothelial cells and, owing to their cargo, they modulate targeted cell behavior. Under hyperglycemic conditions (HGC), endothelial cells changed their properties and became stiffer and less mobile which causes endothelial dysfunction and abnormalities in micro- and macrovascular systems. The aim of this study was to find whether Ect restore mobility and motility of macrovascular endothelial cells under HGC. Uptake of Ect, cell morphology, cytoskeleton organization and membrane stiffness (by atomic force microscopy) were analyzed after the exposure to isolated Ect. To find which cellular pathways were deregulated by HGC and whether Ect could potentially restore gene expression profile, transcriptome analysis was done. We observed that endothelial cells internalized more Ect under normoglycemic conditions (NGC) then HGC. Hyperglycemic cells (HG) were bigger and showed the stiffer surface with denser actin cytoskeleton in comparison to normoglycemic cells. Number of metabolic pathways was influenced under HGC, especially those related to intracellular transport, metabolism and cellular component organization and Ect did not restore HGC impaired cell signaling. Ectosomes cannot reverse this harmful effect of hyperglycemia in endothelial cells, which can have clinical implication in use Ect as therapeutic target in diabetes treatment.

## 1 Introduction

Macrovascular and microvascular complications are one of the leading causes of morbidity and mortality among patients with both type 1 and type 2 diabetes [1]. Hyperglycemia (HG), the main trigger of microangiopathy, leads to severe and life-threatening complications including loss of vision (retinopathy), end-stage renal failure (nephropathy), diabetic cardiomyopathy and peripheral neuropathy [2]. Endothelial cells are among the most vulnerable cells to the harmful effects of high glucose levels, due to their inability to down-regulate glucose uptake during HG [3, 4, 5]. *In vivo* and *in vitro* studies confirmed that HG evokes biochemical and functional changes in endothelium, which triggers endothelial cell dysfunction (ECD) [6]. These changes are manifested as a set of functional alterations, including a reduction in cell proliferation and migration, which results in impaired wound healing as well as enhanced apoptosis [7, 8]. Other important physiological manifestations of endothelial dysfunction are the imbalance between vasodilation and vasoconstriction [9], along with the induction of ischemia and neo-angiogenesis [4, **Error! Bookmark not defined**.]. Functional abnormalities are accompanied by structural alterations in blood vessels, such as the thickening of the basement membrane (BM) [10] and the inner limiting membranes (ILMs) or the reduction of the outer diameter of the capillary tubes. It has been shown that the stiffening in the cells cultured in high glucose conditions (25 mM) is the result of actin cytoskeletal remodeling. These changes are not reversible even after the restoration of normal glucose levels (5 mM) during cell culture [11].

It is possible to distinguish between collective and local cell movements. The first type, collective cell movement, is usually described as cell migration and plays a pivotal role in a variety of physiological and pathological processes such as vasculogenesis, tumor growth, wound healing and tissue regeneration or revascularization [12]. The second type, local cell movement, is related to the structural fluctuations and is characterized by membrane ruffling, withdrawal of retraction fibers, and formation of new lamellae and *filopodia*, leading to local shape changes [13]. These types of motions can be random or guided by specific environmental and internal factors, influencing cell movement or dynamics [14]. In diabetes, endothelial cell migration is significantly tangled, with a consequent negative impact on neovascularization and wound healing [2, 15, 16].

Extracellular vesicles (EVs) are small (up to 1,000 nm in diameter) phospholipid bilayer vesicles, released by cells under physiological and pathological conditions [17, 18]. They are vastly present in most body fluids (blood, amniotic fluid, urine, cerebrospinal fluid, breast milk, saliva, etc.) and their number and characteristics can vary, depending on their origin [17, 19]. EV cargo consists of a variety of macromolecular components, i.e., proteins, lipids, non-coding RNAs and/or DNAs, and is directly related to the biochemical state of the cell of origin [16, 20]. Several studies have shown that EVs have a unique ability to mediate long-distance cell-to-cell communication [16, 21, 22, 23]. Internalization of EVs containing functional and endothelium-ameliorating micro-RNAs by recipient cells can significantly alter their properties as well as metabolic activity and motion, promoting cell migration and proliferation [24]. EV classification is mainly based on their physical properties (size and density) [25] and biological activity, the way they are formed (exocytosis, membrane protrusion and pinching, endocytosis, or apoptosis). Mid-sized EVs, ectosomes (Ect), released from cells, carry an extensive set of proteins specific for the cell of origin [22, 26]. Additionally, Ect can transport DNA, mRNA, and miRNA in a horizontal manner to the target cells, especially from one endothelial cell to another [23]. It has been observed that endothelial progenitor cells (EPC)-derived EVs promote endothelial cell survival, proliferation, and angiogenesis [27]. Microarray and RT-PCR analysis have shown that EPC-derived EVs contain mRNA associated with the *PI3K/AKT* signaling pathway and eNOS. Accumulating evidence reveals a role for all sub-populations of EVs in intercellular communication, with particular relevance to hemostasis and vascular biology [28]. Ect released from endothelial cells to conditioned media, under both physiological and pathological conditions, exhibit cell-stimulating activity [29].

This study aimed to investigate how Ect affect the motile abilities of macrovascular endothelial cells, specifically human umbilical vein endothelial cells (HUVECs), by carrying out wound healing (a scratch assay) in an experimental model of poorly-controlled diabetes. Because wound healing is a physiological process regulated by cell mobility and motility, normo- and hyperglycemic endothelial cell single-cell morphology and membrane stiffness were analyzed before and after the exposure to subsequently isolated Ect. To evaluate the influence of glucose and Ect on HUVECs gene expression, we performed transcriptomic analysis. The effect of uncontrolled diabetes treatment was simulated by temporal (long and short) hyperglycemic (HGC) conditions and compared to normoglycemia (NGC).

## 2 Materials and methods

### 2.1 Experimental overview

In this study, HUVECs were tested under four different culture conditions:

- Long-term normoglycemic conditions (NGC) – HUVECs were cultured in 5 mM glucose after isolation and during experiments (4-5 passages).
- Long-term hyperglycemic conditions (HGC) – HUVECs were cultured in 25 mM glucose after isolation and during experiments (4-5 passages).
- Short-term NGC – HUVECs were cultured in long-term HGC after isolation and transferred to NGC at the beginning of experiments.
- Short-term HGC – HUVECs were cultured in long-term NGC after isolation and transferred to HGC at the beginning of experiments.

During experiments, the cell culture media were enriched with Ect isolated from HUVECs cultured in a long-term NGC or HGC. As a control, we used media without Ect.

### 2.2 HUVEC isolation and culture conditions

HUVECs were isolated from umbilical cords collected during cesarean sections of healthy, adult women, following the bioethical guidelines. Umbilical veins were filled with a mixture of 199 medium (Sigma Aldrich, cat. No. M7653) and 0.25% trypsin (powder, Sigma Aldrich, cat. No. 85450C) with EDTA (380 mg/l, Sigma Aldrich, cat. No. E6758), without Ca^2+^/Mg^2+^ and incubated at 37°C in PBS buffer for 30 min. After incubation, the content of the vein was collected, trypsin was inactivated with FBS (Fetal Bovine Serum, BioWest, cat. No. S181B-500) and the suspension was centrifuged at 250 x *g* for 15 min. Pelleted HUVECs were suspended in NGC or HGC medium for further culturing. HUVECs were cultured in a mixture of 199 medium and SFM (Human Endothelium Serum-Free Medium, GIBCO, cat. No. 11111044), with 10% of FBS (Biowest, cat. No. S181B-500), containing penicillin/ streptomycin (BioReagent, cat. No. P0781) at a concentration of 10,000 units/l (penicillin) and 1 mg/l (streptomycin) and 2 mM L-glutamine (BioWhitaker, Lonza, 200 mM, cat. No. 17-605E), respectively. In HGC, glucose was added to a final concentration of 25 mM (Sigma, cat. No.G7021). All experiments were performed on HUVECs after 4^th^ and 5^th^ passage.

The collection of umbilical cords for this study was approved by the Bioethical Committee of Jagiellonian University in Kraków, Poland with the permission number 122.6120.78.2016 and written informed consent for publication had been obtained from participating patients.

### 2.3 Isolation and characterization of ectosomes

Differential centrifugation method was performed according to a previously published protocol to isolate Ect [30, 31]. After cells reached 80% confluence, the culture medium was replaced with serum-free medium, to avoid contamination with serum EVs. Medium was supplemented with different glucose in concentrations 5 and 25 mM according to the experimental protocol (Tab. 1, Fig 1.A) for 24 h of conditioning. Then, the conditioned media were collected and normoglycemic (Ect NG) and hyperglycemic (Ect HG) were isolated by different centrifugation cycles, according to the protocol.The Ect pellets were used for further examinations.

**Tab. 1.** The 12 combinations of factors used in the study.

| Long-term NGC | Long-term HGC | Short-term NGC | Short-term HGC |
| --- | --- | --- | --- |
| <u>control</u> | <u>control</u> | control | control |
| <u>+ Ect NG</u> | <u>+ Ect NG</u> | <u>+ Ect NG</u> | + Ect NG |
| + Ect HG | <u>+ Ect HG</u> | + Ect HG | + Ect HG |

*control = without Ect, Ect NG = with Ect isolated from HUVECs cultured in long-term NGC, Ect HG = with Ect isolated from HUVECs cultured in long-term HGC. The underlined conditions were applied for single-cell morphology evaluation, stiffness analysis, uptake study, and transcriptomic analysis. NGC – 5 mM glucose, HGC – 25 mM glucose.*

**Fig. 1.**
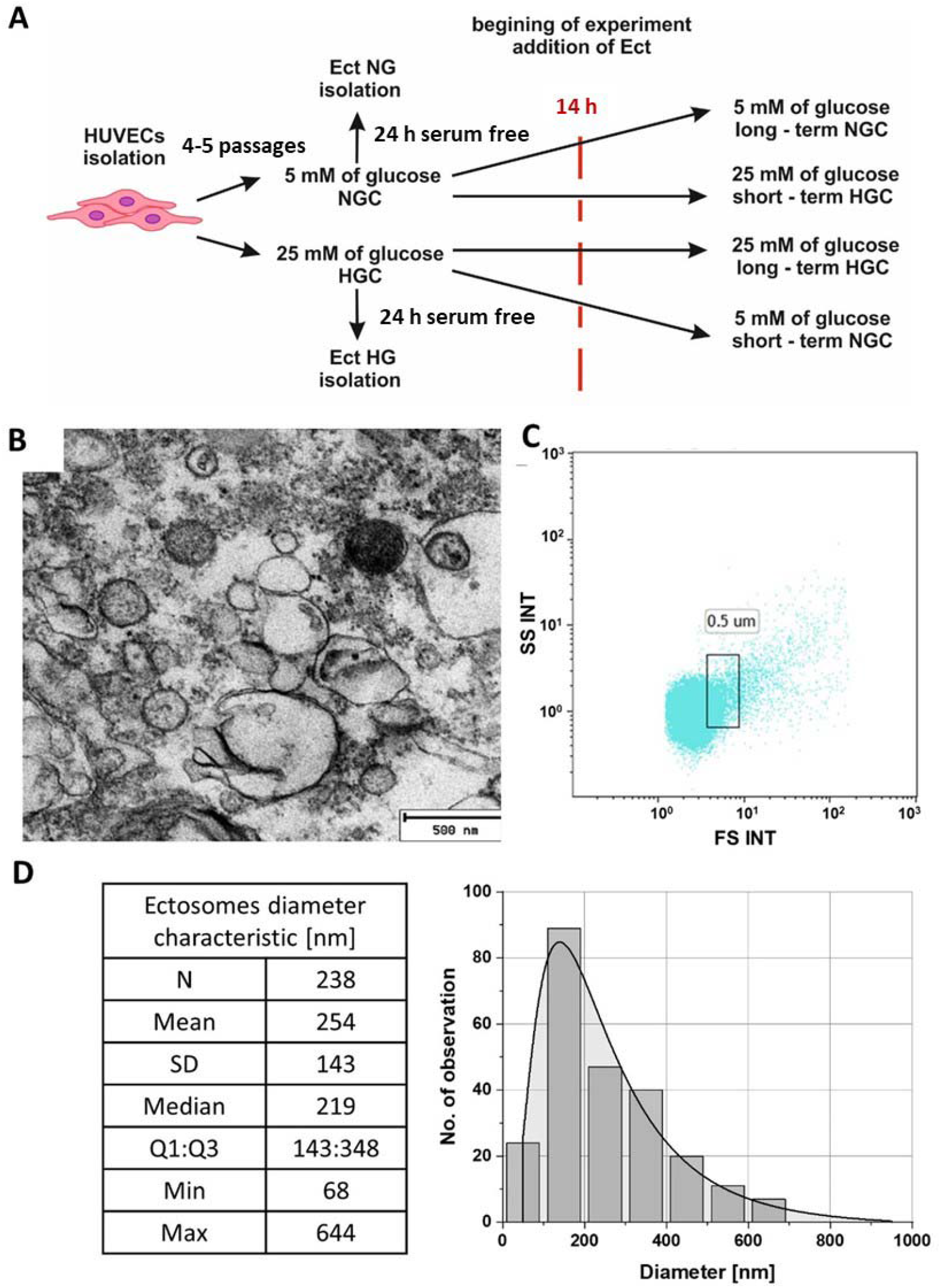
Ectosome characteristics. A – Scheme of the experimental design, B – TEM images of Ect isolated from culture media, C - Flow cytometric analysis of isolated Ect by differential centrifugation (no dye). The gate represents a region of 500 nm calibration beads [**Error! Bookmark not defined**.], D – TEM analysis visualizes the variety of different vesicle-like objects with a diameter of 254 ± 143 nm.

The number and average size of isolated Ect were assessed using flow cytometry (Navios Flow Cytometer, BeckmanCoulter). The Ect analysis region was defined as the upper boundary of the 0.9 μm beads cloud and the threshold right after the cloud from the 0.3 μm calibration beads (*Suppl File*). A representative scatter plot represents the region of data acquisition.

Transmission electron microscopy (TEM) imaging was performed to visualize Ect. Pellets of Ect were processed using a previously described procedure in *Suppl File* [32]. Observation was performed on an electron microscope (JEOL JEM2100HT, Jeol Ltd, Tokyo, Japan) with an accelerating voltage equal to 80 kV.

### 2.4 Cell viability test – Alamar Blue assay

Cell viability test was performed using AlamarBlue dye (AlamarBlue™ Cell Viability Reagent, Thermofisher, cat. No. DAL1100). Three different conditions were tested: long-term HGC with 25 mM and 50 mM glucose concentrations, and long-term NGC (5 nM). Absorbance (570 nm and 600 nm) was measured immediately after media change, after 30 min, then every hour for the first 6 h and after 12 h, 18 h, 24 h, 48 h, 72 h and 96 h [**Error! Bookmark not defined**.].

### 2.5 Scratch assay

A scratch assay was performed using Ibidi inserts (Culture Inserts 2-Well Ibidi, cat. No. 80209) on 12-well plates. Cells were seeded on the inserts at a density of 44,000 cells/cm^2^. Cells were cultured for 24 h in 70 μl of standard culture medium. After 24 h, inserts were removed, cells were washed twice with PBS and placed in serum-free media with glucose in concentrations 5 mM or 25 mM and previously isolated Ect NG or Ect HG, respectively (Table 1).

Microscopic images of the cells were recorded at time 0 (t = 0 h), immediately after the removal of the inserts, and after 14 h of incubation in experimental conditions (t = 14 h). For every artificially created wound (scratch), five images were taken, and the test was repeated 3 times. A confluence index (CI) was chosen as a parameter to assess cell migration. CI is defined as a percentage of area covered by cells after 14 h of incubation, compared to the area of the scratch at time t = 0 h (Eq. 1).

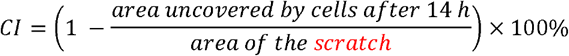

*Eq. 1. Confluence index (Cl) was calculated as a percentage of area uncovered by cells after 14 h of incubation compared to the area of the scratch in time t=0 (right after insert removal)*.

Data were analyzed using diCELLa Scratch PRO software provided by diCELLa – Digital Image Cell Analysis. An example of the CI analysis performed by the software is shown in *Suppl file*.

### 2.6 Single-cell morphology assay

Single-cell morphology observations were performed for selected conditions based on the results of the scratch assay for cells showing the highest CI differences in scratch assay Tab. 1 (underlined conditions). HUVECs were seeded in the 6-well plate at a density of 1,640 cells/cm^2^ to obtain fine, individual cell growth and cultured for 24 h before experiments. Next, cells were placed in serum-free culture media with appropriate glucose levels and previously isolated Ect NG or Ect HG. Cell pictures were recorded for the next 14 h using the Zeiss Axio Observer inverted microscope at 10x magnification (Carl Zeiss Microscopy GmbH, Germany) for further morphology analysis.

For analysis, cell shape was defined by manual contour marking in the recorded images, and cell perimeter was translated into binary stack masks and then analyzed with the “Particle analysis” tool of the FIJI software [33]. Three basic morphological parameters were analyzed: area [μm^2^], circularity [-], and solidity [-] (Fig. 3). The experiments were repeated 3 times, with 38 to 51 individual cells used for further analysis. A detailed description of image analysis protocol is presented in *Suppl file*.

### 2.7 Characterization of cell stiffness using nanoindentation measurements

Stiffness measurements were performed on HUVECs seeded directly on glass slides at a density of 30,000 cells/cm^2^. After 24 h, cell culture media were replaced with cell culture media with the appropriate glucose level and containing previously isolated Ect NG or Ect HG. After 14 h of incubation, media were removed and exchanged for HANKS’s balanced salt solution (Sigma Aldrich, cat. No. H9394), with an addition of glucose (final concentration 5 mM for long - and short-term NGC or 25 mM for long and short-term HGC) and 1% FBS, for the measurements.

Nanoindentation measurements were performed in triplicate using NanoWizard 3 (JPK Instruments AG, Germany) AFM microscope with BioCell™ head (JPK Instruments). The colloidal probe tip (R = 2,25 mm), mounted on V-shaped silicon nitride cantilever, with a spring constant of 0.01 N/m (Novascan, USA) was used for the tests. Data were collected at a speed of 0.8 μm/s and a maximum force of 1 nN. HUVEC samples were placed in the analytical chamber and incubated at 37°C. Data were collected from three sites on each sample from an area of 8 μm x 8 μm, which was divided into 100 (10 × 10) smaller regions. One measurement was performed in each region. To evaluate elastic modulus of HUVECs on the cortex part and deeper cytoplasmic part of cell layers the Hertz model was applied to the indentation curves, using the protocol described previously [**Error! Bookmark not defined**., 34].

### 2.8 Confocal microscopy Ect cellular uptake study

Ect were labeled with PKH26 dye (PKH26 Red Fluorescent Cell Linker Kit for General Cell Membrane Labeling, Sigma Aldrich, cat. No. PKH26GL) as previously described [35]. After staining, the pellets containing PKH26-labeled EVs were resuspended in 1⍰ml of cell culture medium.

For confocal examinations of uptake, HUVECs were cultured on glass slides. After 24 h, cell culture media were replaced with fresh culture media, with appropriate glucose level, containing previously stained Ect NG and Ect HG. After 14 h of incubation, cells were washed three times with PBS and fixed with 3.7% formalin solution in PBS. Cells were stained with phalloidin (Alexa Fluor™ 647 Phalloidin, Invitrogen, cat. No. A22287) to visualize the actin cytoskeleton. Cellular uptake of endothelial EVs was observed and recorded using Zeiss LSM 710 confocal laser microscope with an oil-immersion Plan-Apochromat 40x NA 1.4 objective (Carl Zeiss Microscopy GmbH, Germany), and 633 (phalloidin) and 561⍰nm lasers (PKH26). Samples were prepared in triplicates, and for each sample, five images were taken. To calculate the Ect uptake [%] in each image, the percentage of cells with visible internalized Ect was calculated. Images for long-term NGC and long-term HGC (without Ect) were analyzed in the same way to estimate the nonspecific background fluorescence.

### 2.9 Transcriptomic analysis

Transcriptomic analysis was performed for six previously chosen conditions (Tab. 1). HUVECs were incubated with Ect for 14 h, and subsequently, RNA was isolated from HUVECs using mirVana™ miRNA Isolation Kit (cat. No. AM1560, Ambion) according to the manufacturer’s protocol. Prior to library preparation, RNA samples were evaluated for their integrity, and their concentrations were measured using an Agilent 2100 Bioanalyzer (Agilent) with RNA 6000 Nano Kit (cat. No. 5067-1511, Agilent). Libraries for six samples were prepared using Ion AmpliSeq™ Transcriptome Human Gene Expression Panel (cat. No. A26327, Ion Torrent) that covers over 20,000 human RefSeq genes in a single assay. Libraries were prepared according to the manufacturer’s protocol using 80 ng of total RNA as the input material. Libraries were sequenced on an Ion Proton (Ion Torrent) machine using Ion PI Hi-Q Sequencing 200 chemistry (cat. No. A26433, Ion Torrent) and Ion PI Chip v3 (cat. No. A26771, Ion Torrent). RNA-seq data are available in the Jagiellonian University Repository - RUJ (https://ruj.uj.edu.pl/xmlui/handle/item/176979).

Quality control (QC) checking on raw sequence data was performed using FastQC software (version 0.11.08) [36]. Based on the QC report, low-quality sequences (reads with quality less than Q20, and shorter than 36 bases) and adapters were filtered using trimmomatic software (version 0.39) [37]. Mapping and quantification of clean reads to the Homo sapiens genome hg38 (version 97) were performed using Kallisto (version 0.46.0) pseudo-alignment [38]. TPM (Transcripts Per Kilobase Million) were used to identify differentially expressed transcripts [39]. Transcripts that were not expressed (TPM = 0) in more than three samples were filtered out. Log2FC was calculated and filtered based on following criteria: log2 FC >2 for up-regulated transcripts; and log2 FC < −2 for down-regulated transcripts. Based on the obtained data, the proprietary Python script was used to create a list of statistically significant differentially expressed genes.

Statistically enriched pathways were detected using g:Profiler (GO biological process) [40,41]. Generic Enrichment Map and hsapiens.GO:BP.name.gmt file from the g:Profiler website that contains the gene sets corresponding to GO biological processes, were used as input for visualization using Cytoscape, EnrichmentMap and AutoAnnotate software [42].

### 2.10 Statistical analysis

OriginPro 2016 software (OriginLab Corporation, Northampton, USA) was used for statistical analyses and plots design. The normality test (Shapiro-Wilk test) was performed to assess data distribution. The statistical test was chosen based on the results of the normality test. For further analysis, one-way ANOVA or Kruskal-Wallis test (where normality was rejected) was applied.

To assess the results of the scratch assay and the cellular uptake of Ect, mean values of analyzed parameters were calculated, and measurement uncertainties were expressed using standard deviation. The differences between mean values were analyzed by one-way ANOVA. Differences between subgroups were tested with Tukey’s *post hoc* test; p < 0.05 was chosen for the statistical significance level.

**3** In the analysis of single-cell morphology and cell stiffness, the Kruskal-Wallis test was used to show the measurement uncertainties. The difference between groups was tested with Dunn’s *post hoc* and presented on as mean differences between groups; statistical significance was set at p < 0.05.

## 3 **Results**

### 3.1 Size and concentration

The mean diameter of the ectosomes estimated by TEM was 254 ± 143 nm (Fig. 1B, 1D). The diversity of Ect shape and cargo was presented as different electron density of isolated particles, which corresponds to the particle’s cargo. Additionally, diversity in Ect size was confirmed in flow cytometry analyses. The average concentration of Ect used for experiments was calculated and equal to 1,097 ± 261 particles/μl (Fig. 1C).

### 3.2 Cell viability tests

Cell viability test (AlamarBlue) was performed to assess whether HGC influence cell viability and welfare, which could be linked to altered cell migration or the ability of cells to carry out wound healing processes. There were no statistically significant differences between the analyzed groups [**Error! Bookmark not defined**.].

### 3.3 Scratch assay

Results of the wound healing assay are shown in Fig. 2. Comparison of the control with experimental conditions (first row) showed that there were no statistically significant differences between the CI values. In the presence of Ect NG (second row), CI was increased in HUVECs cultured in both long- and short-term NGC compared to those cultured in long-term HGC (23.25 ± 7.77% vs 10.99 ± 4.45%; p = 9.69*10^−4^ and 26.58 ± 6.02% vs 10.99 ± 4.45%; p = 1.4*10^−4^, respectively) and the short-term HGC group (23.25 ± 7.77% vs 13.72 ± 5.63%; p = 0.01 and 26.58 ± 6.02% vs 13.72 ± 5.63%; p = 9.3*10^−4^, respectively). In the presence of Ect HG (third row), CI in long- and short-term NGC was significantly higher than that of cells tested in long-term HGC (22.18 ± 7.16% vs 13.58 ± 7.67%; p = 0.05 and 22.22 ± 7.69% vs 13.58 ± 7.67%; p = 0.05, respectively).

**Fig. 2.**
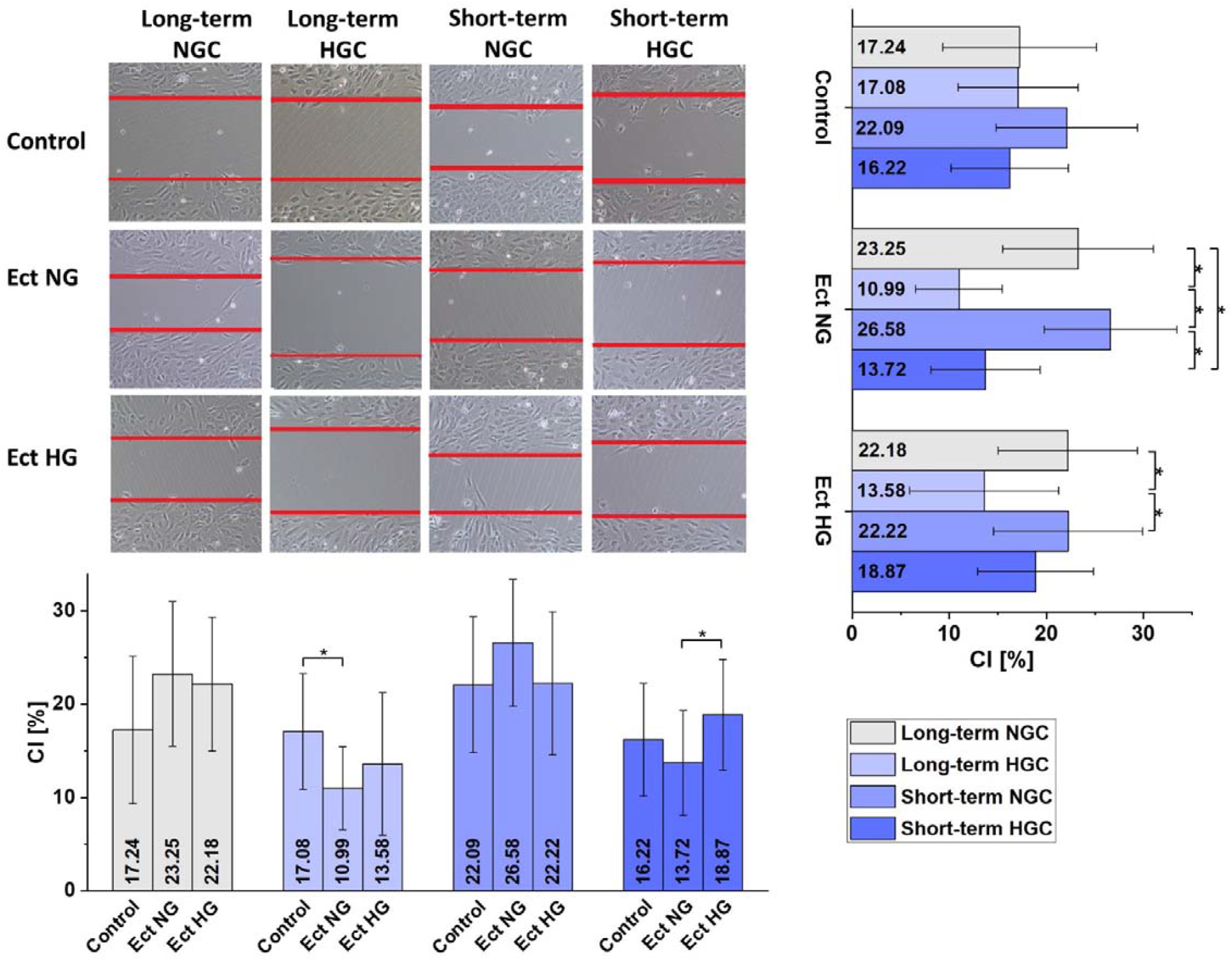
Tabularized results of the wound healing assays under 12 different conditions. Results presented in four columns represent four different cell growth conditions: long- and short-term NGC and HGC. The three rows represent additional factors (Ect NG - Ect isolated from the cells growing in long-term NGC, Ect HG - Ect isolated from the cells growing in long-term HGC) added to cells growing in each condition during the wound assay, and the control group (first row, no additional factors). Images show the exemplary microscopic images of the wounds, created by the cells growing in a given condition, with a given additional factor, after 14 h of incubation. The last column/row shows the graphs of calculated CI values after t = 14 h, for a set of growth conditions, with a stable additional factor (or CI values for a set of additional factors, with a stable growth condition). Data are presented as mean values with the standard deviation (whiskers). One-Way ANOVA test was used. Differences between subgroups were tested with Tukey’s post hoc test and are marked with an asterisk.

Analysis of the differences between four tested growth conditions showed that for both long-term NGC (first column) and short-term NGC (third column) culture conditions, there were no statistically significant differences in the CI values between control cells and cells tested in the presence of either Ect NG or HG. However, for both long-and short-term HGC growth conditions, the differences in cell behavior were more distinguishable. For long-term HGC growth conditions (second column), in the presence of Ect NG, cell migration was impaired compared to the control conditions, without Ect (10.99 ± 4.45% vs 17.08 ± 6.19%; p = 0.04, respectively). For these growth conditions, there were no significant differences in CI between control cells and the cells tested in the presence of Ect HG. For short-term HGC growth conditions (fourth column), Ect HG appeared to stimulate cell migration more efficiently then Ect NG (18.87 ± 7.95% vs 13.72 ± 5.63%; p = 0.03, respectively). Despite this, there were no statistically significant differences between short-term HGC control and long-term NGC and HGC control groups.

### 3.4 Single-cell morphology assay

The results of the single-cell morphology assay are shown in Fig. 3. All three analyzed parameters varied highly between the tested conditions. Between two control groups, long-term NCG and long-term HCG, without the addition of Ect, statistically significant differences in all parameters were observed. Area analysis showed that in comparison to long-term NGC, cells cultured in long-term HGC had a greater area (2,007 ± 1,097 μm^2^ vs 2,778 ± 2,010 μm^2^; p = 6.26*10^−6^, respectively), were more elongated (circularity, 0.68 ± 0.15 vs 0.59 ± 0.19; p = 6.45*10^−13^) and had more irregular boundaries (solidity, 0.87 ± 0.09 vs 0.83 ± 0.12; p = 2.18*10^−11^).

**Fig. 3.**
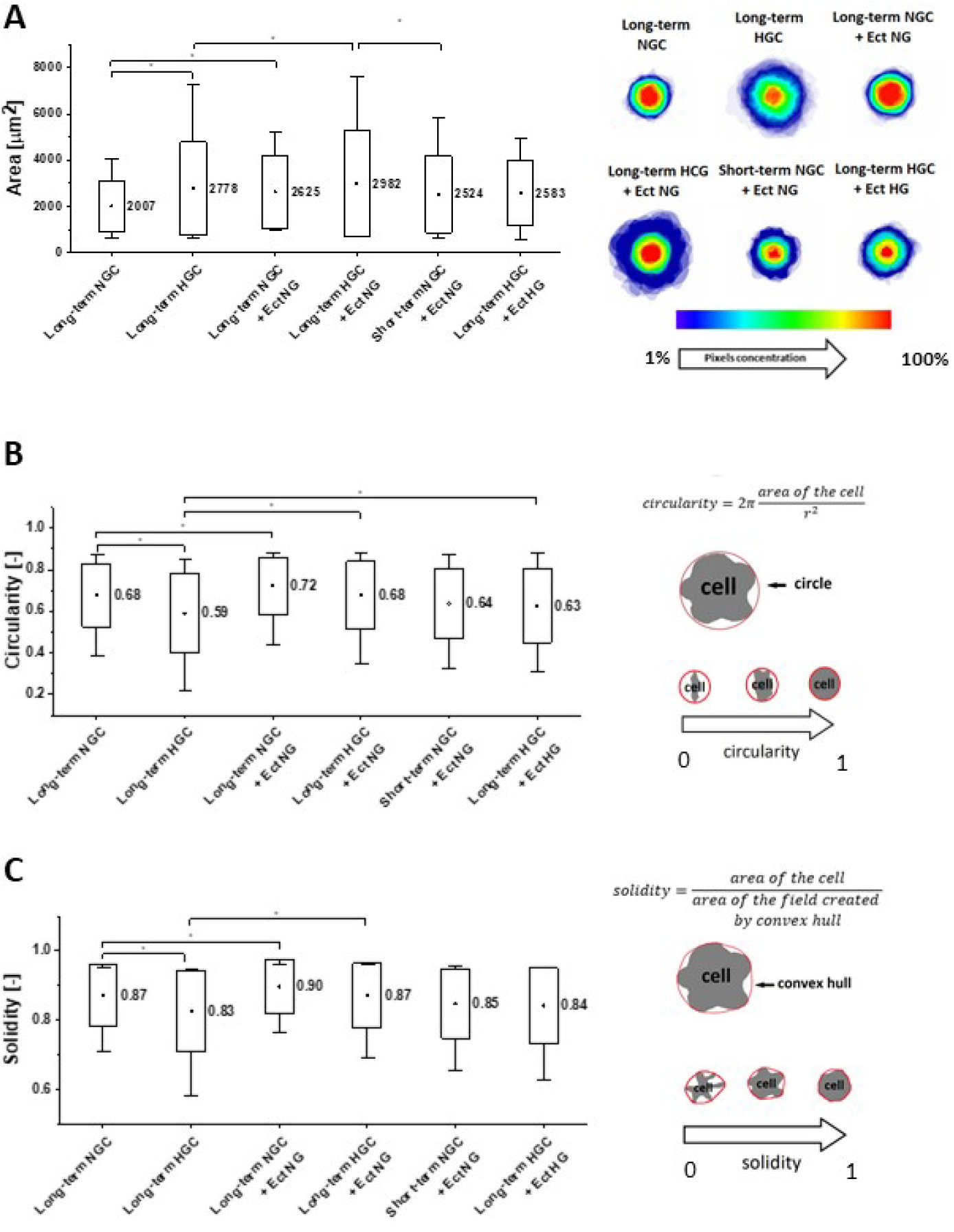
Single-cell morphology assay results. Data are presented as mean values (quad) with median value (line) and the standard deviation (box) and 5-95 percentile values (whiskers). The analysis was performed using the Kruskal-Wallis test. Differences between subgroups tested with Dunn’s post hoc test are marked with an asterisk. A – Graphical presentation of the results of cell area analysis and the stacked images representing cell shape changes. B – Graphical presentation of the results of cell circularity analysis and description of the calculation method. The value 1 represents a perfectly circular shape, while the value 0 represents an infinite line. C – Graphical presentation of the results of cell solidity analysis and description of the calculation method. Solidity is defined as the area of the cell divided by the convex area (area of the field created by convex hull) and was calculated according to the equation. This parameter is equal to one for the shape with no protrusions or irregularities, and zero for the shape with many thin insets [43]. * - p-value <0.05

In the presence of Ect NG, long-term NGC control cells had greater area (2,625 ± 1,574 μm^2^ vs. 2,007 ± 1,097 μm^2^, p = 0.000919, respectively), circularity (0.72 ± 0.14 vs 0.68 ± 0.15; p = 0.02, respectively) and solidity (0.90 ± 0.08 vs. 0.87 ± 0.09; p = 0.03, respectively). For long-term HGC in the presence of Ect HG, when compared with HGC control only, the increase in circularity was observed (0.59 ± 0.19 vs 0.63 ± 0.18; p = 0.05, respectively). For the same cells in the presence of Ect NG, the increase in area (2,982 ± 2,282 μm^2^ vs 2,778 ± 2,010 μm^2^; p = 0.03, respectively), circularity (0.68 ± 0.10 vs 0.59 ± 0.19; p = 1.23*10^−11^, respectively) and solidity (0.87 ± 0.09 vs 0.83 ± 0.12; p = 1.29*10^−10^, respectively) was observed. In the presence of Ect NG, cells cultured in the short-term NGC had a smaller area in comparison with the long-term HGC (2,524 ± 1,657 vs 2,982 ± 2,281; p = 1.22*10^−^4), no differences in circularity and solidity were observed. Additional analysis of differences between groups is available in *Suppl file*.

### 3.5 Cell stiffness in nanoindentation tests

During the nanoindentation measurements, the signal was generated from different layers of a cell. Response from the surface contains information about the membrane and cortex of the cell, while a response from deeper layers is mostly influenced by the cytoskeleton and the nucleus. The results of the nanoindentation measurements are shown in terms of the stiffness of the surface of the cells (Fig. 4) and the deeper layer of the cells (Fig. 5).

**Fig. 4.**
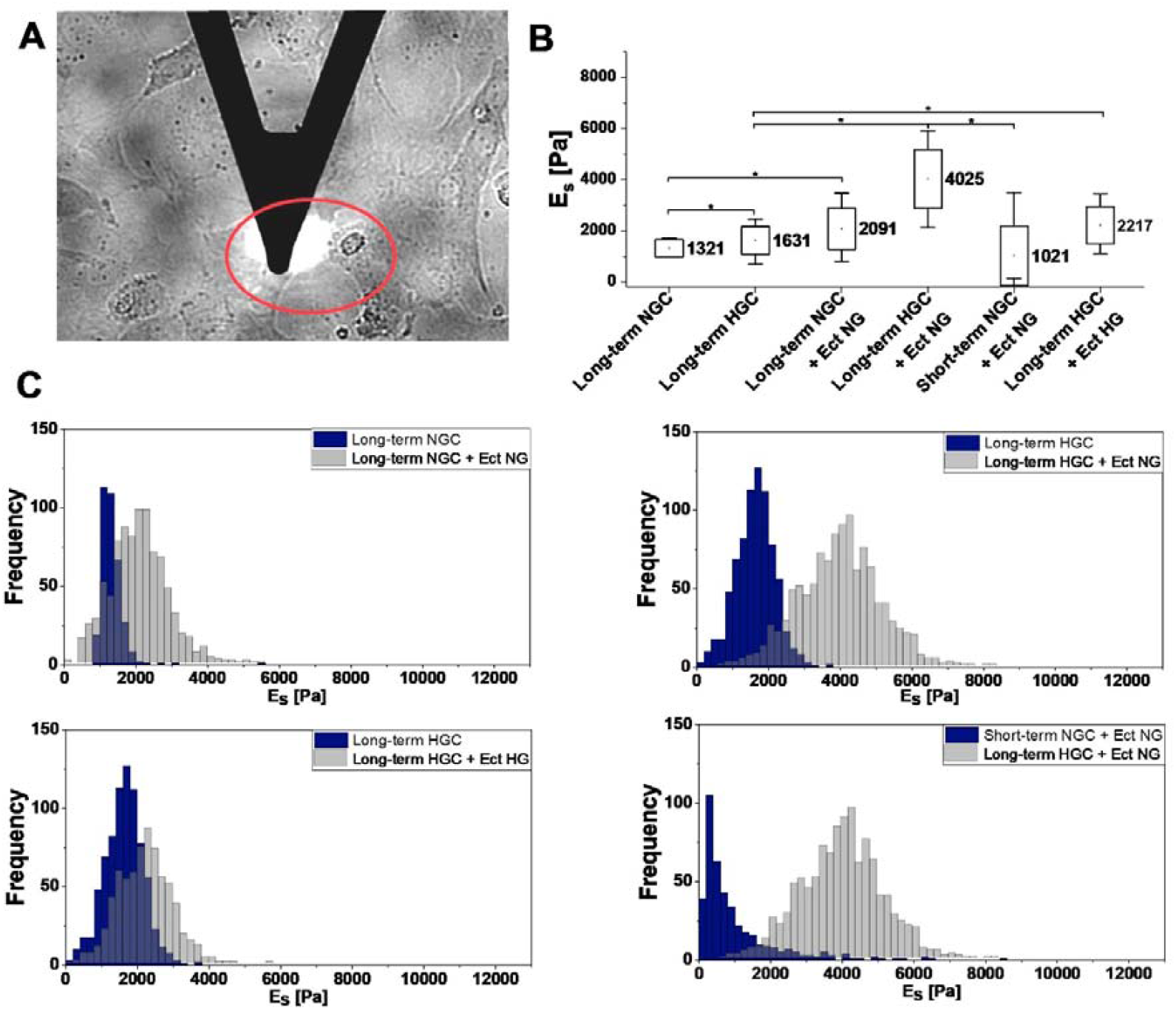
Nanoindentation test results for surface stiffness. A – Image of AFM probe on HUVEC cells. B – Graphical presentation of the mean values of surface stiffness. Data are presented as mean values (quad) with median value (line) and the standard deviation (box) and 5-95 percentile values (whiskers). The analysis was performed using the Kruskal-Wallis test. Differences between subgroups tested with the Dunn’s post hoc test are marked with an asterisk. C – Histograms of elastic modulus. * - p-value <0.05

**Fig. 5.**
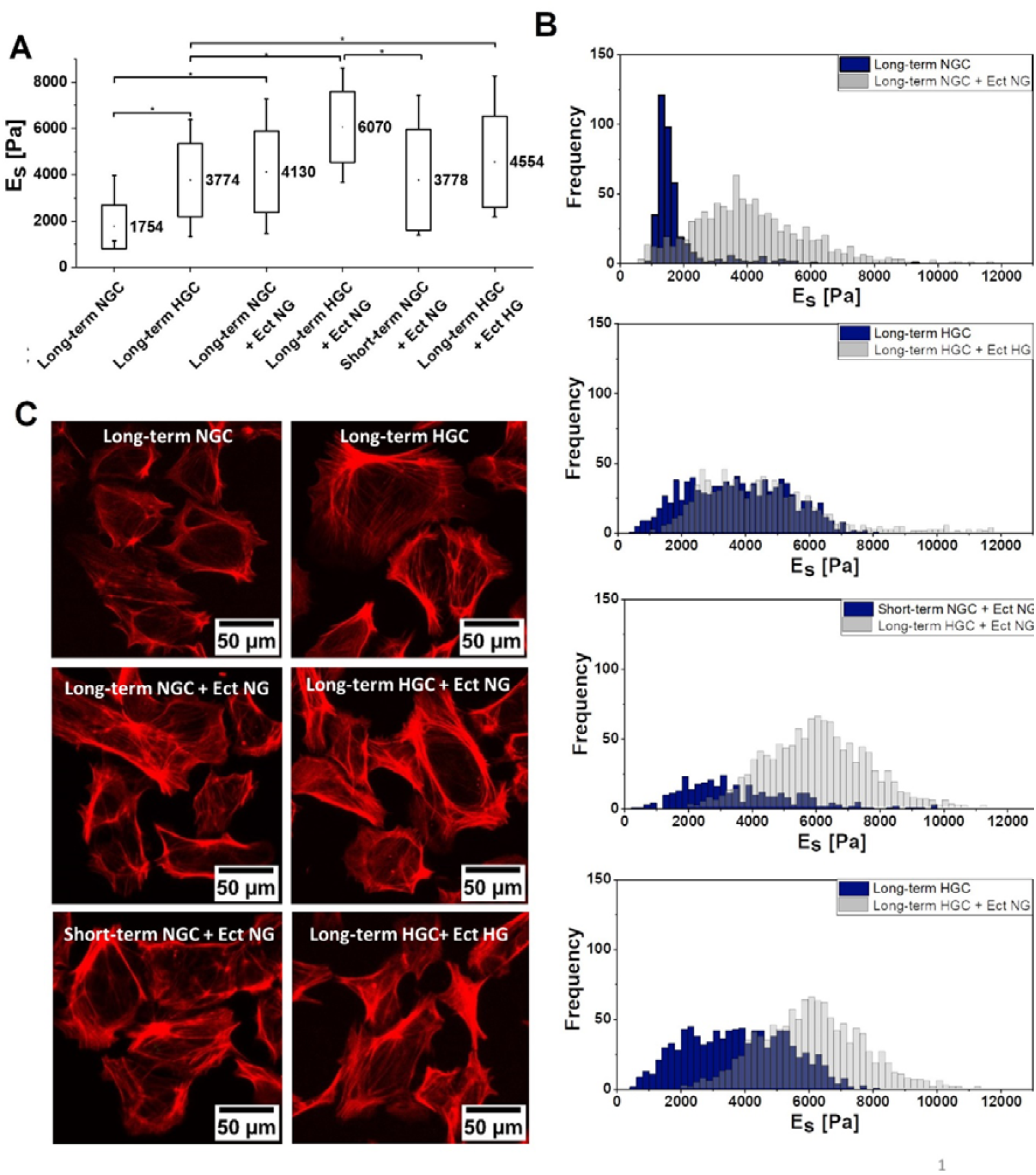
Nanoindentation test results for deeper layer stiffness. A – Graphical presentation of the mean values of a deeper layer stiffness. Data are presented as mean values (quad) with median value (line) and the standard deviation (box) and 5-95 percentage values (whiskers). The analysis was performed using the Kruskal-Wallis test. Differences between subgroups tested with the Dunn’s post hoc test are marked with an asterisk. B. Histograms of elastic modulus. C – Confocal microscopy images of HUVECs cytoskeleton. The cytoskeleton was visualized with phalloidin staining. * - p-value <0.05

Overall, there was a considerable variation in stiffness between the cells in the six previously chosen conditions. For both layers, we observed the same tendency in cell stiffness. Stiffness of cells cultured in long-term NGC was lower than that of cells cultured in long-term HGC both, on the surface and in deeper layers (1,312 ± 344 Pa vs 1,631 ± 542 Pa; p < 0.001 and 1,753 ± 932 Pa vs 3,774 ± 1,579 Pa; p < 0.001, respectively).

In the presence of Ect NG, cell stiffness was increased in comparison to long-term NGC control cells in both surface and the deeper layers (2,091 ± 808 Pa vs 1,312 ± 344 Pa; p < 0.001 and 4,130 ± 1,763 Pa vs 1,753 ± 932 Pa; p < 0.001, respectively). Long-term HGC cells in the presence of Ect HG also had increased stiffness both on the surface and in the deeper layers compared to HGC control (2,217 ± 735 Pa vs 1,631 ± 542 Pa; p < 0.001 and 4,554 ± 1967 Pa vs 3,774 ± 1,579 Pa; p < 0.001, respectively). Similarly, for the cells cultured in HGC in the presence of Ect NG, in comparison to the HGC, increase in stiffness on the surface, and in the deeper layers was observed (3,890 ± 1,053 Pa vs 1,631 ± 542 Pa; p < 0.001 and 6,070 ± 1,530 Pa vs 3,774 ± 1,579 Pa; p < 0.001, respectively). In the presence of Ect NG, cells cultured in short-term NGC in comparison to long-term HGC were characterized by decreased stiffness both on the surface (1,021 ± 1,155 Pa vs 4,025 ± 1,052 Pa; p < 0.001, respectively) and in deeper layers (6,070 ± 1,155 Pa vs 3,778 ± 2,177 Pa; p < 0.001, respectively). Additional analysis of differences between is available in *Suppl file*.

### 3.6 Cellular uptake of Ect

Microscopy analysis confirmed that both Ect NG and Ect HG are internalized by HUVEC cells (Fig. 6). The highest number of cells with internalized Ect was observed for long-term NGC cells incubated with Ect NG (35.21 ± 10.39%) and it was significantly higher than the internalization of Ect NG in long-term HGC cells (22.32 ± 9.40%, p = 0.002), and short-term NGC cells (24.00 ± 8.16%, p = 0.01) as well as Ect HG internalization of long-term HG cells (20.88 ± 10.32%, p = 7.94*10^−4^).

**Fig. 6.**
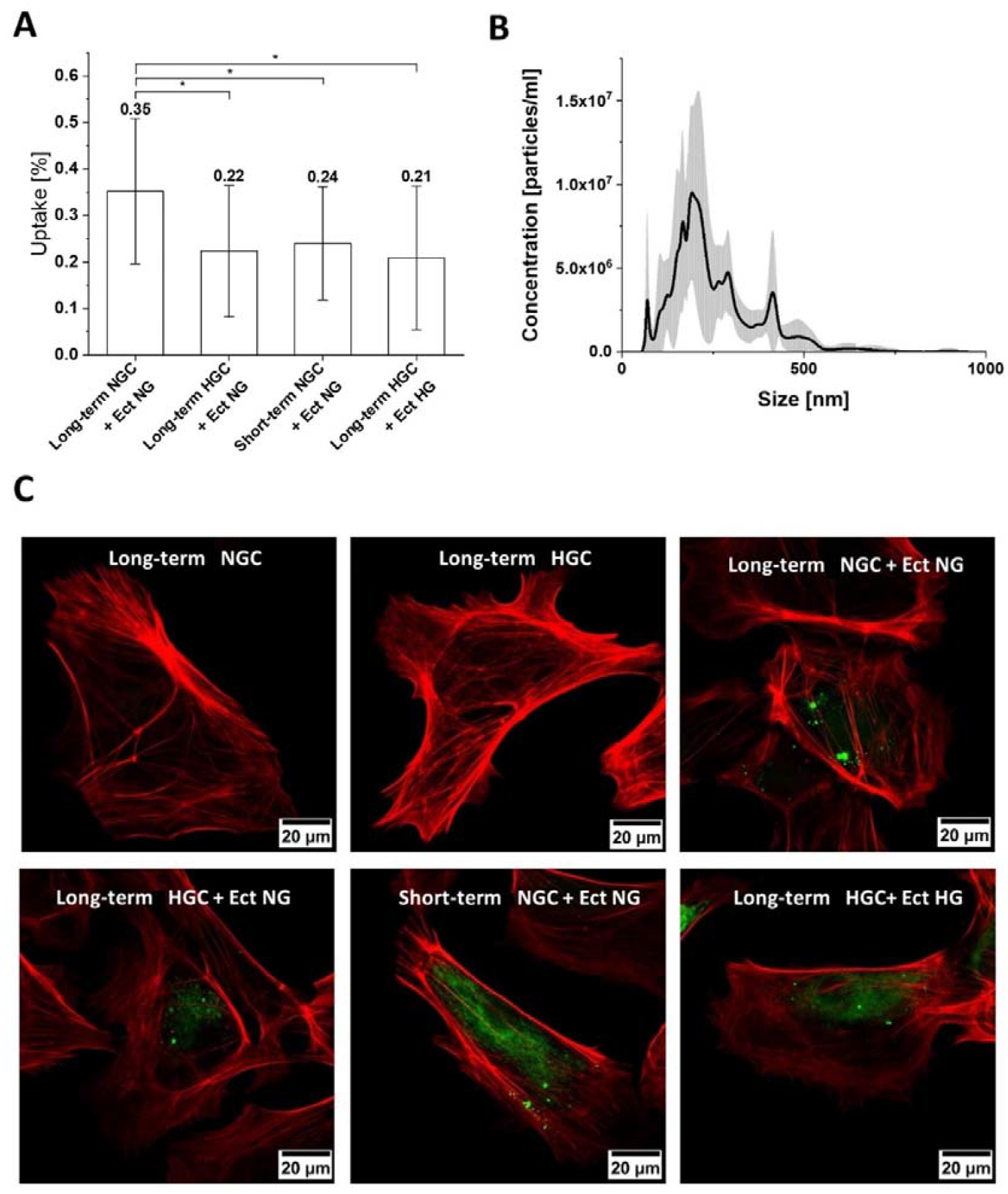
A. Graphical presentation of the Ect cellular uptake for six previously chosen conditions. Results were calculated as percentage of cells with visible internalized Ect. Data are presented as mean values with the standard deviation (whiskers). One-Way ANOVA test was performed. Differences between subgroups were tested with the Tukey’s post hoc test and are marked with an asterisk. B – An example of NTA results of the Ect NG concentration and size distribution measurement. C – Confocal microscopy images of HUVEC cells with internalized ectosomes. Cytoskeleton (red) was visualized with phalloidin staining and ectosomes (green) with lipophilic dye PKH 26.

### 3.7 Transcriptomic analysis

RNA-Seq is a powerful tool that can be used to compare transcripts or gene expression and find up- and down-regulated gens. In this study, we used a bioinformatic pipeline based on systems biology analysis, such as gene ontology, to identify critical biological processes related to the function of differentially expressed genes.

The purpose of the first step of bioinformatics analysis was the identification of expression levels between pairs of samples representing two experimental conditions: a) short-term NGC + Ect NG vs long-term HGC + Ect NG; b) long-term NGC + Ect NG vs long-term NGC; c) long-term HGC + Ect HG vs long-term NGC + Ect NG; d) long-term HGC vs long-term NGC; e) long-term HGC + Ect HG vs long-term HGC and f) long-term HGC + Ect NG vs long-term HGC (Table 2). It was observed that for three pairs: a, b and c the differences were small (123, 146 and 139 genes were respectively identified as differentially expressed). For pairs: d, e and f significant differences were observed (1,049, 1,317 and 1,378 genes were respectively identified as differentially expressed). This (d, e, f) paired gene list was selected for pathway enrichment analysis.

**Tab. 2.** The number of differentially expressed genes identified between two experimental conditions.

| **Analyzed sample** | **Control sample** | **Number of differentially expressed genes** |
| --- | --- | --- |
| short-term NGC + Ect NG | long-term HGC + Ect NG | 123 |
| long-term NGC + Ect NG | long-term NGC | 146 |
| long-term HGC + Ect HG | long-term NGC + Ect NG | 139 |
| long-term HGC | long-term NGC | 1,049 |
| long-term HGC + Ect HG | long-term HGC | 1,317 |
| long-term HGC + Ect NG | long-term HGC | 1,378 |

Pathways that are significantly represented and genes overlapping between pathways in an analyzed samples genes list were identified. Data were visualized by grouping genes representing similar GO biological processes. As a result of this operation, enrichment maps were obtained with clusters of pathways representing the most important biological subjects (Fig. 7.). We found the signaling pathways that regulate metabolic processes (GO:0019222), intracellular transport and localization (GO:0046907), or cellular response to stress (GO:0033554) were significantly affected by long-term HGC alone and by long-term HG with NG and HG Ect presence.

**Fig. 7.**
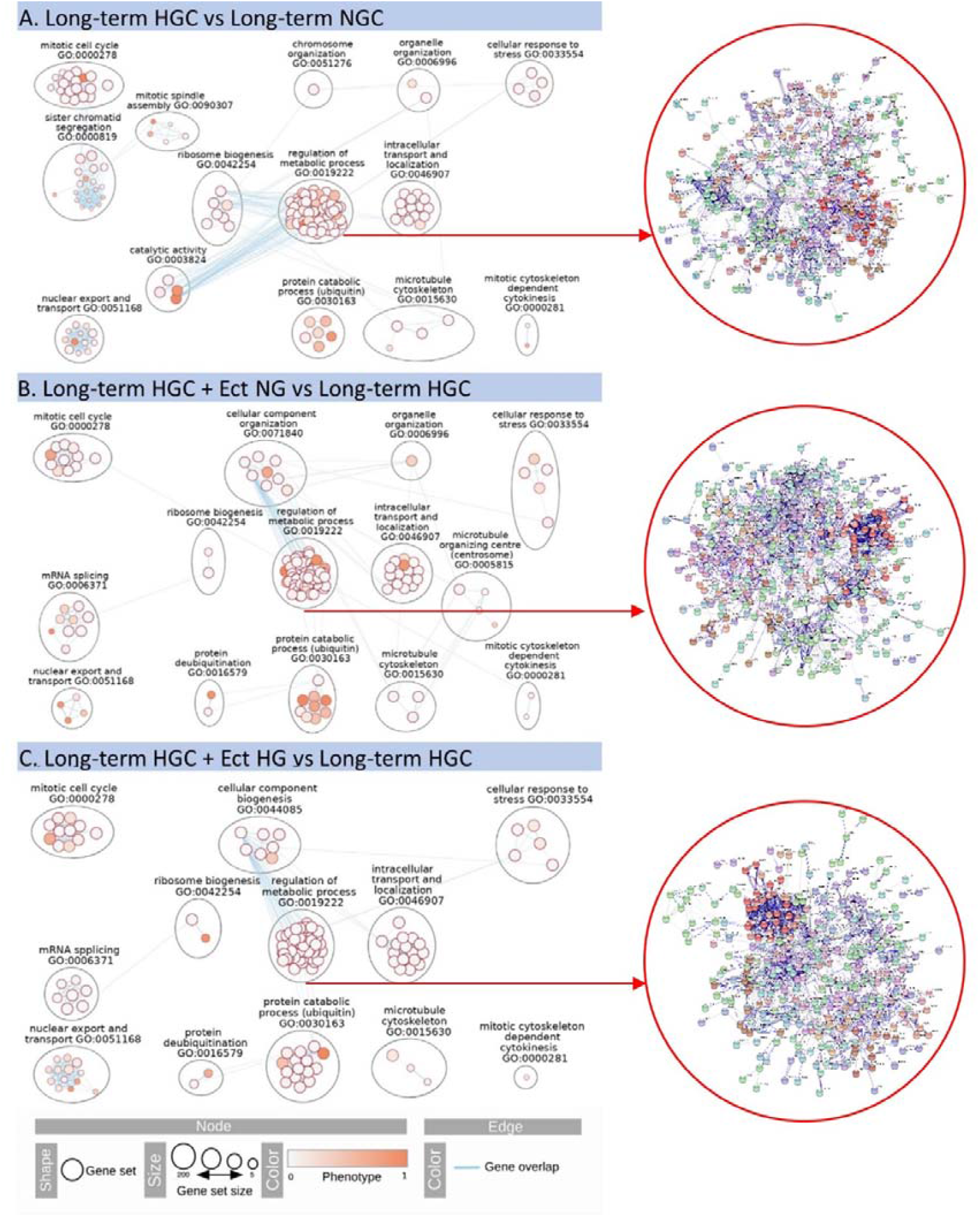
Enrichment map. Created with parameters FDR Q value < 0.01, and Jaccard Index (similarity coefficient) = 0.25. Nodes represent pathways. Edges represent the number of genes which overlap between two pathways.

## 4 Discussion

In our study, we assessed the influence of ectosomes applied on endothelial cells (HUVECs) cultured in diabetes-mimicking conditions and tested Ect ability to improve wound healing within endothelial monolayer. We investigated the role of hyperglycemia in the presence of normo- and hyperglycemic ectosomes on HUVEC morphology as well as their ability to contribute to wound healing. Additionally, we analyzed cell stiffness and single-cell morphology upon Ect uptake under the same conditions. After we confirmed that ectosomes are internalized by HUVEC cells, we evaluated the influence of Ect and glucose level on gene expression in HUVEC cells.

The most significant finding in our study is that the change in the cell culture conditions from HGC to NGC reversed the deteriorating effect of HG on endothelial cells. Thus, we provided evidence that impaired functions and aberrant structure of endothelial cell exposed to high glucose can be reverted upon normalization of glucose levels. The most intriguing observation was that the endothelial cells exposed chronically to high glucose and then transferred to normal glucose gained the highest wound-healing capability, provided they were exposed to ectosomes (Fig. 2). In our study, we used a scratch assay as a model of wound process. This model assumes that a collective cell migration represents the multistep process observed during wound healing in multicellular tissues. However, the wound healing is a process involving a series of biological events taking place leading to scar formation. In such process, not only cell migration contributes, but also cell motility, including lamellipodia formation and cytoskeleton remodeling and cell shape [44]. Cell circularity decreased after long-term HGC to rise significantly after exposure on normoglycemic Ect, similar to solidity (Fig 3. B, C). It means that cells became more flat and regular after Ect uptake. To compare with a scratch assay, cells in the same conditions loss their migratory potential (Fig. 2). Accordingly, the highest value of CI was observed for cells cultured in short-term NGC in the presence of Ect NG (26.58 ± 6.82%). Moreover, we observed that cells cultured in long-term NGC, in the presence of Ect NG were characterized by the second highest CI (23.26 ± 5.15%) - Ect improve wound healing by endothelium under the physiologic glucose conditions. When we compared these results with a significantly lower value of CI for cells cultured in long-term HGC, even in the presence of Ect NG or Ect HG (10.99 ± 4.45% and 13.58 ± 7.67%, respectively), we concluded that HG decreases cell migration and negatively affects the wound healing process. This may be due to changes in the ectosome cargo and changes in cell surface molecules during hyperglycemic cell culture conditions. If we assume that ectosomes are messenger particles, it can be supposed that cells, by releasing Ect, may influence the behavior of surrounding cells in paracrine manner.

Our data on cell level support the clinical doctrine on beneficial effects of glycemia normalization in diabetes. Based on these results, we propose that not only culture conditions but also the fraction of EVs, namely ectosomes (Ect), which can stimulate cell migration and change endothelial cell behavior, significantly influences cell migration. The test results showed that high glucose did not influence tested cell viability [**Error! Bookmark not defined**.]. Thus, we propose that the surrounding environment, i.e., HGC and the Ect NG or Ect HG, influences cell behavior without affecting cell viability and resulted from the impact of Ect, provided that glucose levels were reversed to normal.

Previously it was shown that EVs can influence the wound healing process. Jansen et al. [24], in an *in vitro* wound healing assay, proved that the migration of human coronary artery endothelial cells (HCAEC) is enhanced in the presence of endothelial cell-derived microparticles (cultured in NGC). They also observed that microvesicles released from cells, cultured in HGC conditions, lost their stimulating potential. In another study by Zhang et al. [45], it was found that HUVECs migration is enhanced in the presence of EVs derived from human-induced pluripotent stem cell-derived mesenchymal stem cells (hiPSC-MSC). In the study on normoglycemic and diabetic wound fibroblasts migration, enhanced migration *in vitro* was observed in both normal and diabetic wound fibroblasts after treatment with MSC-derived EVs [46]. Additionally, the study on exosomes (Ex) derived from HUVECs collected from women suffering from gestational diabetes mellitus (GDM) (Ex_GDM_), and healthy pregnant women (Ex_HC_) confirmed that Ex can influence wound healing processes. It has also been observed that in the case of HUVECs from healthy donors, wound recovery was more efficient than for GDM HUVECs. Moreover, in this study, it has been shown that Ex_HC_ can stimulate GDM HUVECs and increase the efficiency of the wound healing process, while Ex_GMD_ has the opposite effect, i.e., wound healing efficiency is decreased [47]. In our study, we observed that the most responsive were Long-term NGC cells exposed to Ect NG. The differences in the results in Sáez’s *et al*. and our study results probably from the variations in the EV population used. Exosomes differ from Ect in terms of protein, lipids content and as we can assume in different internalization pathway [25, 26]. Exosomes are internalizes via clathrin dependent mechanism but Ect may be internalized by fusion, which may produce direct effect on the target cell, including migratory behavior.

In our study, we observed that endothelial cell viability was not affected by hyperglycemia. These data are consistent with the study by Apostolova et al. [48], where using the same assay they confirmed that HUVEC viability is not decreased in the presence of glucose at concentrations of 5 mM, 25 mM, 50 mM and 100 mM in 96 h; however in the samples measured after 96 h of incubation in the presence of glucose (50 mM and 100 mM), cell viability was decreased. Opposite results were obtained by Karbach *at al*. [49]. Using trypan blue staining method, they showed that glucose (30 mM) gradually decreased HUVECs viability, calculated as the number of living cells during a five-day experiment.

It is known that Ect contains biologically active molecules (e.g., proteins, miRNA, etc.), which can influence the cell activity, including cell migration. It has been proven that macrophage foam cell-derived EVs stimulate vascular smooth muscle cell (VSMC) and HUVEC migration, probably through the regulation of the actin cytoskeleton and focal adhesion pathways [50]. Another study revealed that HCAEC-derived EVs promote vascular endothelium repair by delivering functional miR-126 into the recipient cells [24]. Additionally, it has been proven that the EMP-mediated miR-126-induced-cell repair process is impaired in endothelial cells under HGC [2424]. This observed attenuation in endothelial healing could be the effect of changes in the cell membrane composition caused by HGC, which in turn can influence membrane transport and cell signaling. Previous studies proved that hyperglycemia reduces glycocalyx volume of the endothelial cells [51]. Moreover, the high glucose in a time-dependent manner decreased the expression of CD59 and CD55 in HUVECs, which is probably linked to the ECD, by the activation of the complement pathway [52].

The results of the single-cell morphology assay performed in this study showed that cell culture conditions and Ect can influence cell shape parameters. The average cell area was larger for cells cultured in HGC, and the area was increased in the presence of Ect in both NGC and HCG cell culture media. We observed that in long-term NGC, cells in the presence of Ect NG caused the increase in their diameter, which was correlated with the increase in circularity and solidity. Changes in the culture conditions from NGC to HGC reduced the cell area but did not appear to influence cell solidity and circularity. The same tendency was observed for the long-term HGC group in the presence of Ect NG. Only circularity and solidity were altered compared to the same cells without Ect. To the best of our knowledge, we demonstrated for the first time that high glucose concentration and the presence of Ect influence cell shape. This phenomenon could be linked to the osmolarity of cell culture media or the impact of the presence of Ect on the cell osmolarity.

The cell stiffness analysis performed under the same conditions as the single-cell morphology assay, for both surface and deeper cell layers, revealed the same changes. We observed that in HGC, cell stiffness was generally increased, which could be linked to the results of our cell shape analysis with one exception. Additionally, we observed that both in the presence of Ect NG and Ect HG, cell stiffness was increased. Surprisingly, the increase in stiffness was higher for Ect NG than that of Ect HG.

The influence of high glucose concentrations on cell stiffness has been known for a long time. It has been proven that cell stiffness is increased in HGC, and it is linked to the alterations in the cytoskeleton. Moreover, restoring NGC does not reverse the changes caused by high glucose levels and cell stiffness remains elevated in relation to NGC [11]. Authors of the above-mentioned study named this behavior as the ‘stiffness memory’ and concluded that such memory is responsible for persistent and not reversed cell stiffness [11]. In another study, the relationship among cell stiffness, NO production, and ECD caused by the inflammatory factor, tumor necrosis factor (TNF)-α was tested [53]. They showed that TNF-α-induced changes in the elasticity of the endothelium were negatively correlated with NO production and associated with the reorganization of the actin cytoskeleton. In our study, cell stiffness, both surface and deeper layers, increased significantly (p<0.5) after Ect NG exposure. This observation suggest that Ect uptake, which is decreased under hyperglycemic conditions, induced changes in cytoskeleton organization (Fig. 5C). Actin cytoskeleton forms a strong cortex, which stabilize cell surface and contribute in cell migration observed as lower IC under long-term HGC with Ect NG in a scratch assay.

Interestingly, constant as well as varying high glucose levels change membrane permeability of endothelial cells. These alterations can be linked to the redistribution of the F-actin bundles, which form the cytoskeleton [54]. We hypothesized that such changes are affected by the inhibition of eNOS expression and NO synthesis caused by high glucose levels. Not only the stiffness of a single-cell but also more complex structures such as blood vessels are affected by glucose concentrations. It was observed that the stiffness of the ILMs of, e.g., retina blood vessels, is increased in the samples collected from the patients suffering from diabetes [55]. Additionally, it was reported that exosomes can influence arterial stiffness. Chen et al. [56] analyzed the levels of circulating microparticles (MP) in blood taken from patients with type 2 diabetes and correlated these results with the brachial-ankle pulse wave velocity (baPWV). They found that the MP level of blood correlates with baPWV values. Based on logistic regression analysis, they concluded that age, HbA1C, total cholesterol, blood pressure, annexin-V^+^ MP, leukocyte-derived MP, and endothelial cell-derived MP were significant risk factors of severe arterial stiffness in diabetes. Additional multivariate analysis showed that endothelial cell-derived MPs are the most significant and independent risk factor for severe arterial stiffness in diabetes.

For transcriptomic analysis, we designed the experimental protocol to consider physiological conditions of uncontrolled diabetes and we used Ect isolated from long-term culture conditions to simulate the effect of uncontrolled diabetes, in which long periods of high glucose levels are alternated with short periods of physiological glucose levels after taking antidiabetic drugs. The stimulation time with Ect was constant (14 h) which was calculated to observe the effect of ectosomes on cells (transcription and metabolic pathways activation), to avoid the effects of other external conditions.

Pathway enrichment analysis showed that important signaling pathways were modified in long-term HGC (alone and with NG or HG Ect), including pathways that affect the regulation of metabolic processes (GO:0019222), intracellular transport and localization (GO:0046907), and cellular response to stress (GO:0033554). This result supports the hypothesis that Ect are a very strong regulation and communication factor. Their effect is detectable even when using invasive external factors that affect whole cells. There are some evidences that endothelial cell-derived Ect are able to impair endothelial cell function by enhancing superoxide formation and increasing oxidative stress in these cells [57]. Moreover, detailed analysis of miRNAs signature of Ect isolated from plasma of patients suffering from diabetes has shown that these extracellular vesicles contain factors which act multi-directional or sometimes in the opposite way: pro- and anti-angiogenic [23].

Our analysis indicated that Ect has an effect on HUVECs metabolism: regulation of metabolic process GO:0019222, protein catabolic process (ubiquitin) GO: 0030163 or protein deubiquitination GO:0016579. A number of papers described the key role of extracellular vesicles in metabolic diseases by transporting enzymes and fatty acids [58,59]. Presence of insulin-signaling proteins was also shown in EVs (including phospho-p70S6K, phospho-S6RP, phospho-GSK3β, phospho-Akt, phospho-insulin receptor (IR), phospho-IRS1 (Ser312), tyr-phospho-IRS1, phospho-IGF-1R, leptin receptor, and FGF21) [3]. Additionally in patients with T2DM, there is an increased presence of EVs containing clusters of differentiation like: CD31, CD41, CD51, CD61 or CD11b [59,60]. These molecules have a great influence on metabolism, which has been demonstrated by reducing their levels using dietary interventions [60,61].

Another important cellular processes were influenced in HUVECs by Ect: cell division, growth and intracellular organization: cellular component biogenesis GO:0044085, ribosome biogenesis GO:0042254, organelle organization GO:0006696, intracellular transport and localization GO:0046907, mitotic cytoskeleton dependent cytokinesis GO:0000281, microtubule cytoskeleton GO: 0015630. These mechanisms involve molecules transported by Ect, such as growth factors including transforming growth factor beta (TGFβ), segregated agents such as interleukin-1b (IL-1b) [62] or matrix remodeling enzymes like matrix metalloproteinases or aggrecanases, heparanases and hyaluronidases [63].

It has been already proven that extracellular vesicles are internalized by endothelial cells [35]. Although, there are several mechanism responsible for extracellular vesicles internalization all of them requires remodeling of the cell surface and could led to the changes in cell stiffness. Moreover in the pathological conditions, as hyperglycemia, stress fibers could be formed what may additionally increase cell stiffness. On the other hand, processes such as cell division or cell migration requires remodeling of the cytoskeleton, mostly actin remodeling, so dependent of the cell conditions cell stiffness can fluctuate. Thus, cell stiffness is a parameter that can describe cellular activity which is not always related to their function improvement and is rather related to remodeling of the cell membrane and the cytoskeleton.

Finally, we can assume that in our study, endothelial cells cultured in NGC receive signals transmitted by Ect, so the increase in the cell area, solidity, circularity, and wound healing (CI) happens much more significantly and effectively than in HGC. Results from cellular uptake of Ect analysis support this hypothesis. During NGC endothelial cells release Ect which are able to stimulate neighboring cells in a paracrine manner to enhance their migration. During ectosome formation some molecules, including proteins (cytokines), are selectively sorted into the ectosome cargo and this process depends on the current state of the parental cell and blood glucose levels [64]. Because of that, Ect NG and Ect HG may carry the cargo with different protein and miRNA composition. An alternative explanation is changed competence of HUVECs in HGC, the lack of stimulation by Ect HG results from a decrease in HG Ect incorporation by recipient cells. In our research we have confirmed the decline in Ect uptake (Fig. 6).

We observed that Ect NG are most efficiently internalized by cells cultured in long-term NGC. It is likely that, with lower uptake, cells received weaker stimulations from Ect NG. In terms of cells cultured in HGC, neither single-cell morphology or other parameters were significantly altered, nevertheless, not only one but two or more parameters were changed, with or without the accompanying changes in CI and cellular uptake of Ect. It is possible that the HGC may cause cell “insensitiveness” to Ect stimulus, and these changes are observed in functional tests. To understand better the mechanism of endothelial cell response, in vivo angiogenic functions and wound healing functions an animal model should be proposed: type 1 and type 2 diabetic mice (db/db and ob/ob mice). However it is noteworthy, that there are not a good model for T2DM, which reflects human conditions.

## 5 Conclusions

Our results show that glucose level and Ect affect the cell’s ability to perform wound healing. Cells cultured in HGC are efficiently stimulated by Ect when shortly exposed to normal glucose. Cells cultured in HGC show reduced internalization of Ect and appear to be insensitive to stimuli transferred by these particles, unless the glucose levels become normal. This lower Ect internalization could be explained by the alterations in cell surface properties, although more research is needed to confirm this hypothesis. Harmful effect of hyperglycemia can be reversed by changing the culture conditions to NGC. Ect are those factors responsible for the normalization of cellular parameters (as size, stiffness etc.). Intriguingly, the changes resulting from HGC to NGC make the endothelial cells the most effective in cell migration provided Ect were added.

## Acknowledgments

The authors want to express their gratitude to Dr Eng Olga Woźnicka from the Microscopy Laboratory of the Institute of Zoology and Biomedical Research, Jagiellonian University in Kraków, for specimen preparation and TEM imaging, Prof Hubert Huras from the Department of Obstetrics and Perinatology, Faculty of Medicine, Medical College, Jagiellonian University in Kraków for collecting umbilical cords and Prof Francisco J. Enguita from Instituto de Medicina Molecular, Faculdade de Medicina, Universidade de Lisboa for help in the analysis of gene expression results.

