## Supplementary File for "Ectosome effect on endothelial monolayers in hyperglycemic and normoglycemic conditions"

***Supplementary File A***

**Ectosomes isolation procedure - differential centrifugation**

HUVEC cell culture in 80% of confluence were washed and cultured 24 h in serum-free medium to avoid contamination with serum EVs. To release normo- and hyperglycemic ectosomes (Ect NG and Ect HG), glucose in concentration 5 and 25 mM was used. Then, the conditioned media were collected in 50 mL Falcon tubes and the Ect isolation was performed. To remove cell debris, conditioned media were subjected low-speed centrifugation cycles: 400 x *g* for 10 min, 3,100 x *g* for 25 min and 7,000 x *g* for 20 min. After each centrifugation cycle, the supernatant was transferred to the new tube to undergo the next cycle. In the last step, supernatants were centrifuged at 18,000 x *g* for 20 min. The upper part of the supernatant was discarded, and the 1.5 ml of the bottom part of the supernatant was transferred to the Eppendorf tube and centrifuged at the same conditions. Then, Ect were washed three times with 300 μl PBS and the operation was repeated.

**TEM sample preparation**

Ect pellets were fixed with 2.5% glutaraldehyde (Sigma Aldrich, cat. No. G5882) in 0.1 M cacodylic buffer (Sigma Aldrich, cat. No. C0250) and then postfixed in 1% osmium tetroxide solution (for the duration of 1 h). The samples were then dehydrated, by passing through a graded ethanol series, and embedded in epoxy resin PolyBed 812 at 68°C (Polyscience, Inc., cat. No. 08791-500). Ultrathin sections were placed on the 300 mesh copper grids covered with formvar film. Sections were contrasted using uranyl acetate and lead citrate.

**Flow cytometry calibration**

Flow cytometry analysis was performed without staining to evaluate the number of Ect present in every sample. Samples were previously concentrated by differential centrifugation of conditioned media. Before each analysis, flow cytometer was calibrated using Megamix-Plus FSC (Megamix-Plus FSC, Biocytex, cat. No. 7802) to ensure an appropriate level of stability of measurements. The exemplary dot plot from calibration is presented in Fig. A1.

The Ect analysis region was defined as the upper boundary of the 0.9 µm beads cloud and the threshold right after the cloud from the 0.3 µm calibration beads

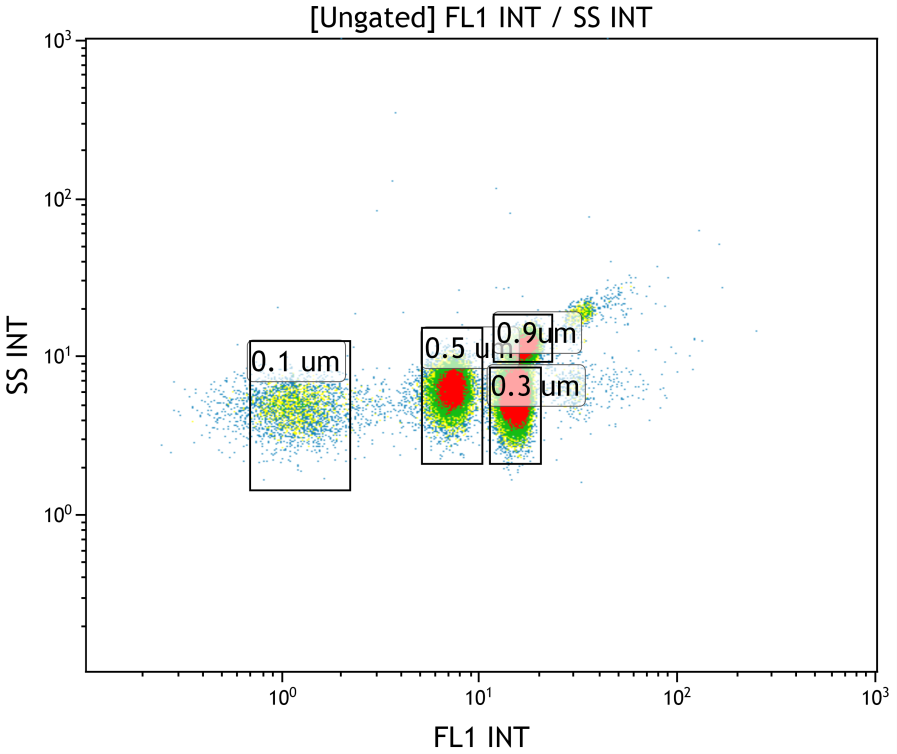

***Fig. A1.*** *Dot plot of the Megamix-Plus FSC calibration beads. Megamix-Plus FSC is a mixture of fluorescent beads with different diameters (0.1 µm, 0.3 µm, 0.5 µm, 0.8 µm). All four groups are well separated.*

**Wound healing assay – image processing**

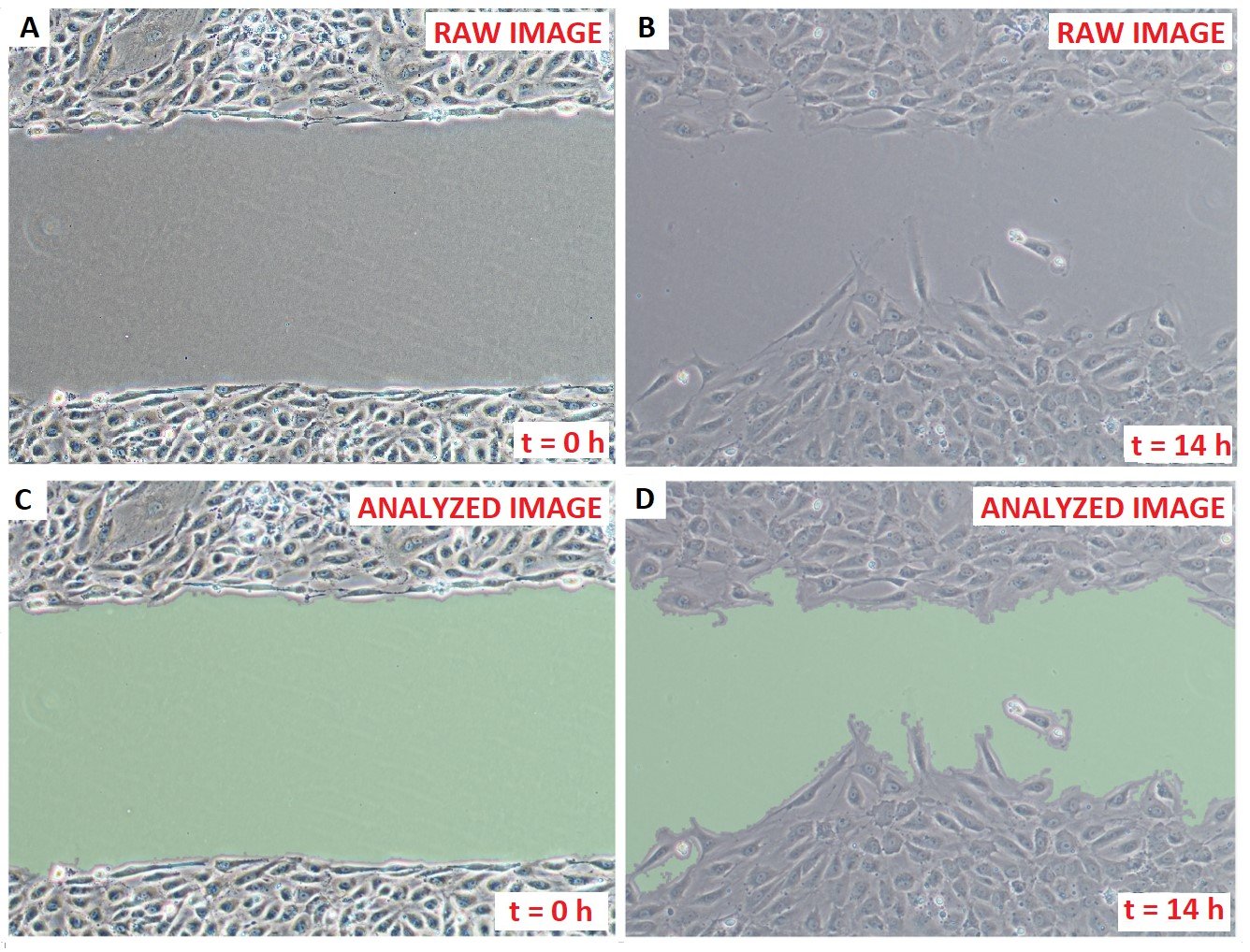

**Fig. A2.** Image analysis of wound healing assay pictures. Raw pictures of HUVECs A – before and B – after the wound healing assay. C,D – Same images were processed by diCELLa Scratch PRO image analyses software. Green areas are recognized by the software as the areas of the wound.

**Single-cell morphology assay**

For the purpose of the study, four neighboring regions of interest were chosen as an area with 10-18 separately growing cells, and the pictures were recorded every 2 h. For further analysis, only cells fulfilling two criteria have been taken:

a) cells must be alive during the whole time of experiment

b) cell must adhere to the surface for at least of 50% of the duration of the experiment.

For a better visualization of the changes in the shape of the cells, stacked images of nine different cells (which were used to calculate area, solidity and circularity), in eight different time points (from t = 0 h to t = 14 h taken every two hours) were created. On the binary images, the center of mass parameter was calculated using the ‘Particle Analysis’ tool. Afterwards, each cell was cut from the image using a square frame (170 x 150 pixels) and positioned in such a way that the center of mass was at the intersection of the diagonals of the square. Next, the stacks were created using all previous images and converted from binary images to heat map images using a LUT (lookup table) tool. In the heat maps, red color represents the highest concentration of pixels in the stack and the blue color represents the lowest concentration of pixels in the stack.

**Atomic force microscopy – Hertz model fitting**

AFM provides a unique way for measuring the nanomechanical properties of biomaterials, i.e., cells or tissues. The elasticity of the sample is determined from the so-called force-distance or force-indentation curves, which reflect the bending of the cantilever during the motion of the piezo-scanner when the AFM probe is pushed against the sample. Standard approach for the analysis of the elasticity data from AFM measurements use the Hertz model [^[1]^, ^[2]^]. According to the Hertz model for a spherical probe, the force $F$ that is required to produce an indentation $\delta$ is given by [^[3]^]:

$$F=kd=\frac{16}{9}E\sqrt{R}\delta^{\frac{3}{2}}$$

***Eq. A1.*** *Force and indentation relationship*.

where:

$k$*– stiffness of cantilever*

$d$ *– deflection of the cantilever*

$E$ *– Young modulus of the sample*

$R$ *– effective radius of the probe*

*δ – indentation*

Usually, the value of $E$ is obtained from the so-called force-distance curves by fitting Eq. A1 to the experimental data. Considering the depth-dependent cell profile, various structural components of the cell have an influence on its mechanical properties [^[4]^]. The nanoindentation data obtained for the studied endothelial cells show the two distinct part that corresponds to the cortex part and the cytoplasmic part of the cell. The cortex part is dominated by the membrane and cross-linked actin filaments located just beneath the plasma membrane. The inner-cytoplasmic counterpart is related to the inner part of the cell and is dominated by nucleus as well as cell cytoskeletal proteins like microtubules and actin microfilaments.

Therefore, we performed a piecewise fit of the data using standard Hertz function from Eq. A1 as visualized in Fig. A2. The value EL corresponds to the apparent elastic modulus for the cortex part and ES to the apparent elastic modulus of the inner-cytoplasmic part. First, a Fit of Eq. A1 was performed for the area close to the maximum retraction of the piezo-scanner (blue area in Fig. A2), for which the force-distance curve is dominated by the contribution from the inner-cytoplasmic part of the cell. This fit was performed using a simple linearized least-square method and is very similar to the procedure described by Sokolov et al. [46]. The fit was performed assuming that the indentation $\delta$is related to the piezo position $Z$ and to the cantilever deflection $d$ by $\delta=Z_{S}^{0}-d-Z$, where $Z_{0}^{S}$ is a fit parameter that corresponds to a “virtual” contact point, i.e., position for which the elastic properties of the substrate are negligible. Next, a similar procedure was performed for the data range near the actual contact point (red/pink range in Fig. A2), where the data is dominated by the contribution from the cortex part of the endothelial cell.

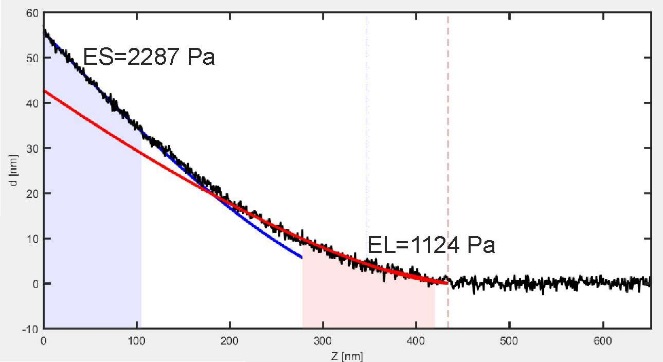

***Fig. A3.*** *Procedure for the determination of the elasticity of the endothelial cell by means of two fits of the Hertz curves. These fits were subsequently performed in the two shaded regions and are marked with blue and red lines. The first (blue) one corresponds to the cytoplasmic part of the cell and represents the stiffest part of the cell. The second one (red part, EL) corresponds to the cortex part of the cell.*

1. [] Radmacher M. (1997) Measuring the elastic properties of biological samples with the AFM. IEEE Eng Med Biol Mag 16:47–57. https://doi.org/10.1109/51.582176.
2. [] Haase K, Pelling AE (2015) Investigating cell mechanics with atomic force microscopy. J R Soc Interface. 12:20140970. https://doi.org/10.1098/rsif.2014.0970.
3. [] Sokolov I, Dokukin ME, Guz N V (2013) Method for quantitative measurements of the elastic modulus of biological cells in AFM indentation experiments. Methods 60:202–213. https://doi.org/10.1016/j.ymeth.2013.03.037.
4. [] Szymonski M, Targosz-Korecka M, Malek-Zietek KE (2015) Nano-mechanical model of endothelial dysfunction for AFM-based diagnostics at the cellular level Pharmacol Rep 67:728–735. https://doi.org/10.1016/j.pharep.2015.05.003.
